# Prescription Opioid Analgesic Use and Mortality in Systemic Lupus Erythematosus

**DOI:** 10.1101/395343

**Authors:** Romy J. Cabacungan, Clifford R. Qualls, Wilmer L Sibbitt, William A. Hayward, James I. Gibb, Selma D. Kettwich, Roderick A. Fields, N. Suzanne Emil, Monthida Fangtham, Arthur D Bankhurst

**Author notes:** Corresponding Author: Wilmer L. Sibbitt, Jr., MD, MSC 10 5550, 5th FL ACC, Department of Internal Medicine, Division of Rheumatology, University of New Mexico Health Sciences Center, Albuquerque, New Mexico, USA 87131, tele.

## Abstract

**Objectives:** This research investigated the prevalence of opioid analgesic use in patients with systemic lupus erythematosus (SLE).

**Methods:** This 5-year prospective cohort study of 275 SLE patients focused on prescription opioid use and 5-year outcome. Associations were determined with univariable regression analysis and then multivariable models were created to determine independent effects on dependent variables

**Results:** Prescription opioid use was common in SLE with 24% using opioid analgesics chronically and 76% not using opioids. Opioid users had a higher rate of tobacco use (p<0.01), cocaine use (p<0.002), mean pain scores (p<0.001), disease activity (SLEDAI-2K) (p<0.001), disease damage (SLICC/ACRDI) (p<0.001), non-adherence to medical therapy (p<0.01), and total deaths at 5 years (opioids: 48.0%, no opioids 19.0%, p<0.001). Logistic regression analysis predicting death revealed opioid use (hazard ratio 2.6, p<0.001) and SLEDAI-2K (1.1, p<0.001) respectively; and opioid use (hazard ratio 2.5, p<0.002), SLEDAI-2K (hazard ratio 1.1, p<0.001), and non-adherence (hazard ratio 1.6, p=0.11), respectively. Multivariable Cox Model analysis estimating probability of death with covariates: opioid use (hazard ratio 2.6, p<0.001) and SLEDAI-2K (hazard ratio 1.1, p<0.001); opioid use (hazard ratios 3.0, p<0.001), and cocaine use (hazard ratio 3.2, p<0.001). The Kaplan-Meir survival analysis revealed a significantly higher probability of death for SLE patients using opioid analgesics.

**Conclusions:** Prescription opioid analgesic use is common in SLE and is associated with markedly increased mortality. Preferably, non-opioid approaches to treat chronic pain should be used in SLE patients.

**Clinical trial registration number:** This was not a clinical trial.

**KEY MESSAGES:** 1. Chronic opioid analgesic use is common in SLE (24%).

2. Opioid use is associated with greater disease severity, tobacco use, non-adherence, and increased mortality.

3. Opioids should be used cautiously in SLE; alternative non-opioid management of pain is recommended.

**ACKNOWLEDGMENTS AND FUNDING INFORMATION:** This work was supported by US National Institutes of Health research grants to Dr. Sibbitt (R01 NS035708) and to the Clinical and Translational Research Center (UL1TR001449).

## Background/Purpose

Systemic lupus erythematosus (SLE) is often characterized by substantial chronic pain that can become a serious long-term management challenge (1-5). As recently reviewed by Borenstein et al and others, opioid analgesics are sometimes incorporated into pain management in order to relieve suffering, avoid acetaminophen and non-steroidal anti-inflammatory drug toxicity, preserve renal function, and reduce corticosteroid use (6-8). However, routine opioid analgesic use for chronic pain can result in dangerous opioid abuse and dependence as recently reviewed by Shipton et al (9). Although there has been important research into the effects of opioid analgesics on the outcomes chronic arthritis as recently reported by lociganic et al and others, the literature for epidemiology of opioid-based chronic pain management in SLE is limited (10,11). In the present study, we report the prevalence of opioid analgesic use in a well-characterized SLE cohort and the comparative 5-year outcomes of SLE patients who were treated or not treated with opioid analgesics for the therapy of chronic pain.

## Methods

This research was approved by the institutional review board (IRB) and was in compliance with the Helsinki Declaration and subsequent revisions. Each subject provided written informed consent. Inclusion criteria included any patient with SLE, age 18-80. Exclusion criteria were age < 18 years, age > 80 years and any patient with an autoimmune diagnosis other than SLE. The study was not an incipient cohort, rather was a cross-sectional study design where the patients were enrolled as they were encountered in rheumatology clinic, consented, information gathered, and then outcomes determined over a 5-year period of observation. The population sample was ethnically diverse and included predominately North American Hispanics (58%) and Caucasian Whites (34%) (Table 1). To be enrolled in the study, all subjects had to be cared for by the university rheumatology practice long-term. Subjects were only enrolled through clinic, and not through in-patient hospital consultations where the patient may have been transferred to the university center from other communities. Because of recruiting through clinic, individuals with extreme life-threatening SLE requiring hospitalization were excluded unless they later came to the clinic for long-term follow-up where they could be enrolled. The diagnosis of SLE was established in each subject using the American College of Rheumatology (ACR) revised criteria for the classification for SLE [12,13]. SLE disease activity was determined with the System Lupus Erythematosus Disease Activity Index 2000 (SLEDAI-2K) and SLE disease injury was measured with the Systemic Lupus Erythematosus Collaborating Clinics/American College of Rheumatology Damage Index (SLICC/ACR-DI) [14,15]. Each of these metrics was further subcategorized into Neuro-SLEDAI-2K consisting of the neurologic components of SLEDAI-2K, and Neuro-SLICC/ACR-DI consisting of the neurologic components of SLICC/ACR-DI (16,17). Neuropsychiatric SLE (NPSLE) was characterized by the ACR nomenclature and case definitions for NPSLE [18].

**Table 1.**
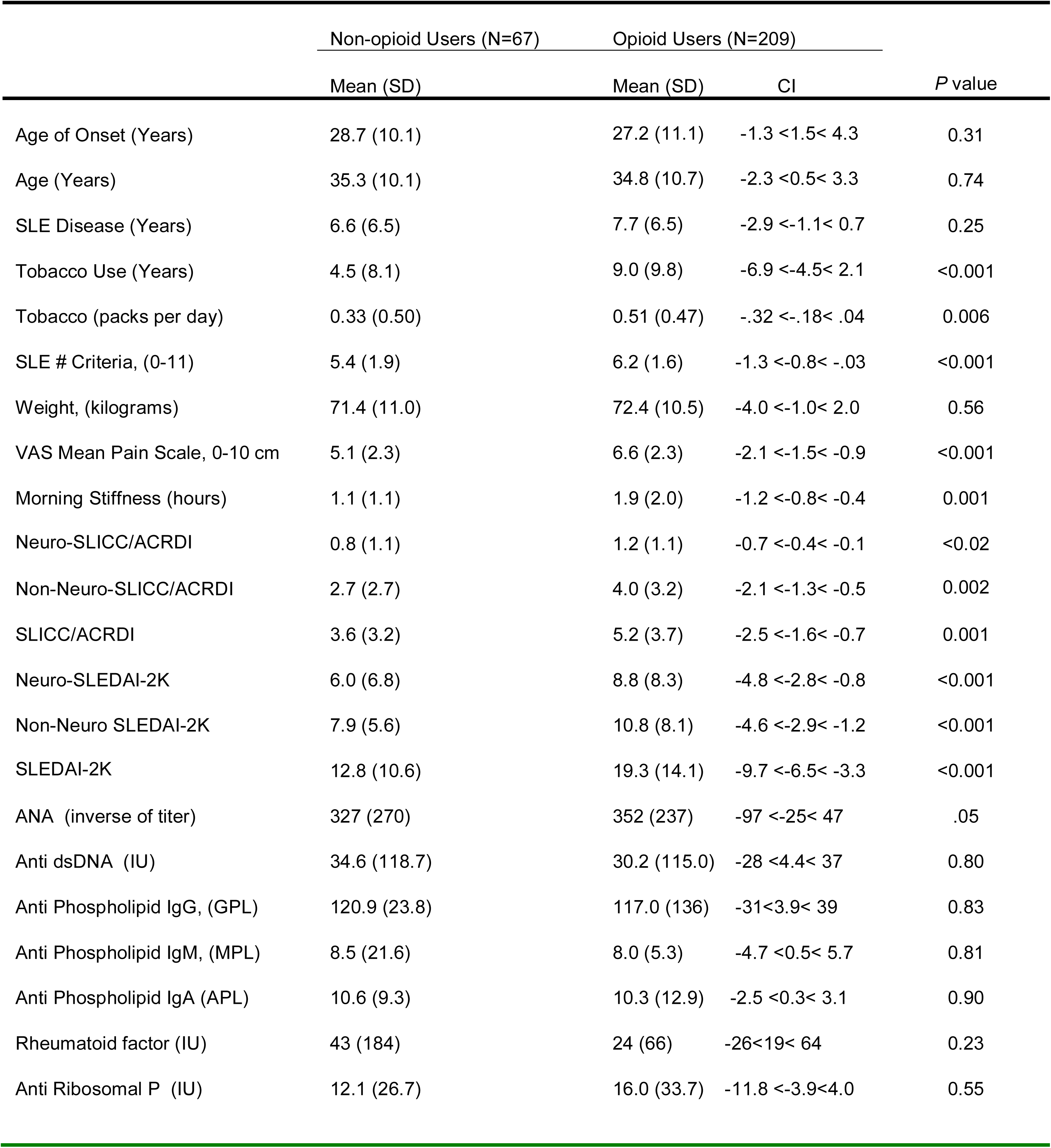
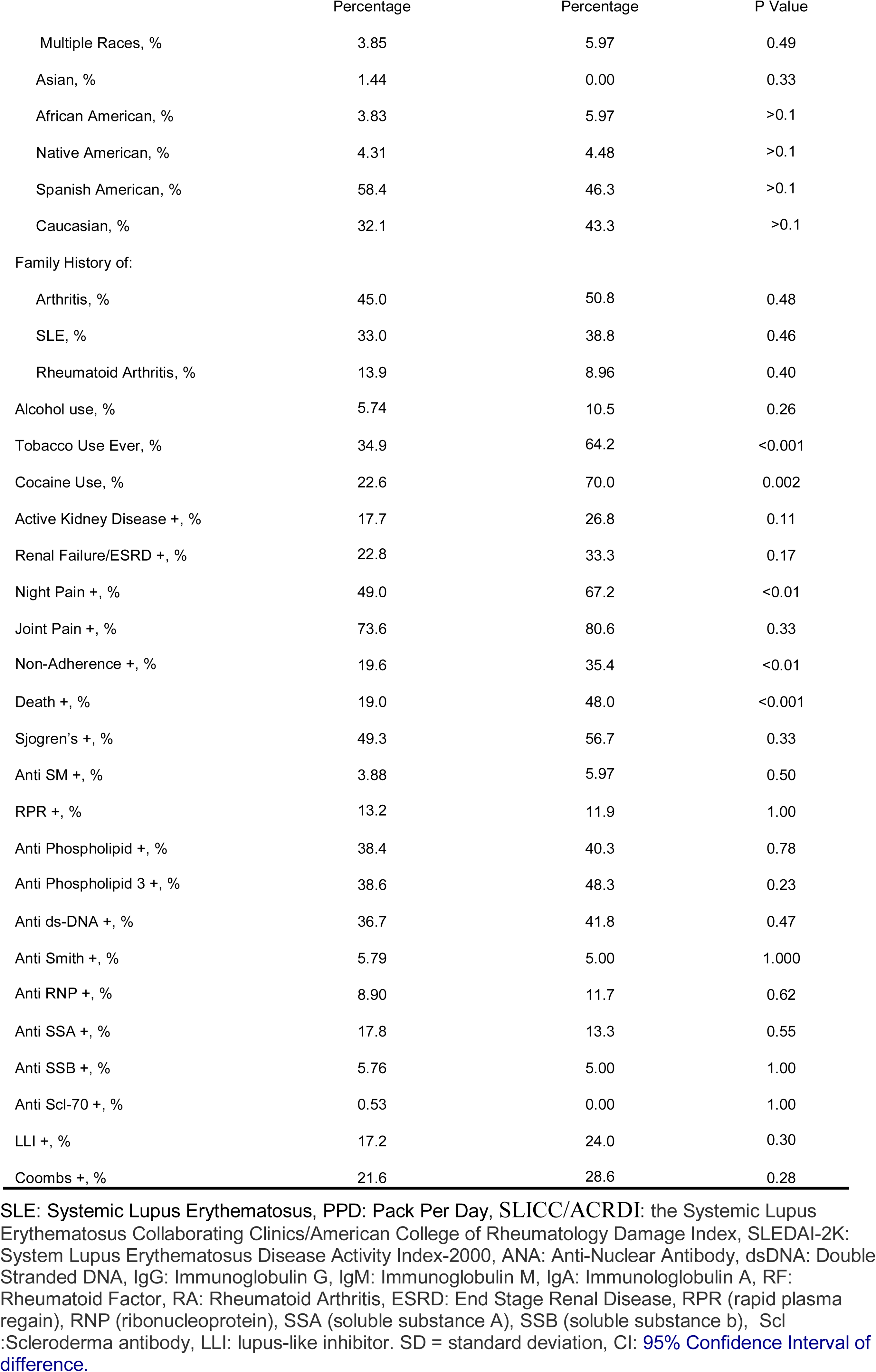
Characteristics of SLE Subjects

Opioid dosage (the study population predominately used oxycodone 5-10 mg tablets) was normalized to 7.5 mg morphine milligram equivalent (MME) (equivalent to 5 mg oxycodone), and was reported as the number of 7.5 mg MME tablets per month (19). Other demographics and outcomes were age of SLE onset (years), age (years), SLE disease duration (years), tobacco use (years), tobacco use (pack per day), tobacco use ever, cocaine use, alcohol use, number of SLE criteria (0-11), weight (kilograms), pain (0-10) visual analogue pain scale (VAS) (20, 21), morning stiffness (hours), SLICC/ACRDI, Neuro-SLICC/ACRDI, Non-Neuro-SLICC/ACRDI, SLEDAI-2K, Neuro-SLEDAI-2K, Non-Neuro-SLEDAI-2K, antinuclear antibody (ANA) titer, anti-DNA antibody (titer), anti-phospholipid antibodies (IgG, IgM, IgA in GPL, MPL, APL units, respectively), rheumatoid factor (IU), antiribosmal P (IU), family history (arthritis, SLE, rheumatoid arthritis), active glomerulonephritis, renal failure/end-stage renal disease, night pain, death, Sjogren’s syndrome, anti-Smith antibodies, anti-RNP antibodies, anti-SSA, SSB antibodies, anti-Scl-70 antibodies, lupus-like inhibitor, direct Coomb’s antibody, false positive syphilis serology (RPR, VRDL), and antiphospholipid antibody (positive/negative).

Non-adherence was defined by admission by the patient that they were not taking their SLE medication, low serum mycophenolate levels, and/or by missing more than 20% of their clinic appointments (a “no-show” - cancelled appointments and hospitalizations were not counted as a “no-show”). Only subjects who could be reasonably followed for 5 years were enrolled, and to ensure this, contact information was obtained from the patient, their relatives, and friends including birth date and social security number so the Social Security Death index could utilized (22). Survival and death of SLE subjects were determined in clinic, by reviewing medical records, following up with telephone calls with each patient and/or the patient’s family and friends, autopsy reports from the medical examiner, chart review, newspaper obituary search, and confirmation of death with the Social Security Death Index for death and benefit files (22). 100% of the 275 subjects were accounted for using these combined methods at the end of the individual 5-year observational period. After collection this prospective 5-year database was then de-identified including removal of all personal information and identifiers. Additional IRB approval was obtained prior to analyzing the de-identified database in relation to opioid analgesic use. Outcomes were determined at 5 years after enrollment in the study.

### Statistical Methods

The major comparison was between opioid users and non-opioid users. Statistical differences between measurement data were determined with Student t-test and between categorical data with Fisher’s exact method; associations were determined initially with univariable regression analysis and afterward multivariable models were created to determine independent effects on dependent variables. Kaplan-Meier survival curves were constructed with regards to important time-to-event outcome variables. Missingness was not an issue due to the use of combined methods to optimize death tracking including strict entry criteria that required local residence, extended family and friends contacts, an individual’s governmental identification, the exclusion of individuals without valid governmental identification, autopsy files from the medical examiner, and the search for missing subjects with examination of medical records, newspaper obituary, and governmental Social Security Death and benefit indexes (22).

## Results

Demographics of the 275 subjects with SLE are shown in Table 1. Opioid analgesic use was common in SLE with 24% using chronic opioid analgesics and 76% not using opioid analgesics. No significant difference was observed in age, age of onset, SLE disease duration, ANA titer, dsDNA antibodies, anti-phospholipid antibodies (IgG, IgM, IgA), rheumatoid factor, anti-ribosomal P antibody, race, family history (arthritis, SLE, rheumatoid arthritis), alcohol use, active kidney disease, end-stage renal disease, joint pain, Sjogren’s syndrome, anti-Smith antibody, RNP, RPR, anti-ds-DNA anti-bodies, anti-SSA and anti-SSB antibody, anti-Scl-70 antibody, lupus-like inhibitor, and direct Coombs antibody in the opioid and non-opioid SLE groups. However, SLE patients who used opioids had statistically signficant increased tobacco use, tobacco smoking duration, packs of cigarettes per day, number of criteria for SLE diagnosis, mean pain by VAS scale, morning stiffness, night pain, SLICC/ACRDI indices (Neuro-, Non-Neuro, and Total), SLEDAI-2K indices (Neuro-, Non-Neuro, and Total), prevalence of cocaine use, non-adherence, and total deaths (Tables 1).

Logistic regression analysis predicting death revealed hazard ratios of 2.59 and 1.06, when comparing opioid use and total SLEDAI-2K respectively, and hazard ratios of 2.48, 1.06, and 1.60, when comparing opioid use, total SLEDAI-2K, and non-adherence respectively (Table 2).

**Table 2.** Logistic Regression Analysis Predicting Death

| Predictor | *B* | Chi-square | *p* | Odds Ratio<br>for Logistic<br>Regression |
| --- | --- | --- | --- | --- |
| Opioids Use | 0.907 | 9.66 | <0.002 | 2.48 |
| Total SLEDAI-2K | 0.062 | 43.60 | <0.0001 | 1.06 |
| Non-Adherence | 0.471 | 2.58 | 0.11 | 1.60 |
| Opioid Use | 0.950 | 10.72 | 0.0011 | 2.59 |
| Total SLEDAI-2K | 0.060 | 44.12 | <0.0001 | 1.06 |

The Univariate Cox Model to estimate for probability of death in SLE patients, which was found to be statistically significant for opioid use and non-adherence, with hazard ratios of 3.22 and 1.81 respectively (Table 3).

**Table 3.** Univariate and Multivariate Cox Models for Estimation of Probability of Death in SLE Patients with Opioid Use

| Variable | Univariate |  | Multivariate |  |
| --- | --- | --- | --- | --- |
|  | Hazard Ratio<br>(95% CI) | P-value | Hazard Ratio<br>(95% CI) | P-value |
| Opioid use | 3.22 (1.89-5.65) | <0.001 | 2.59 (1.46-4.57) | 0.001 |
| Total SLEDAI-2K |  |  | 1.06 (1.04-1.08) | <0.001 |
| Opioid use | 3.22 (1.89-5.65) | <0.001 | 3.28 (2.02-5.34) | <0.001 |
| Alcohol use |  |  | 0.46 (0.11-1.88) | 0.28 |
| Opioid use | 3.22 (1.89-5.65) | <0.001 | 2.99 (1.68-5.33) | <0.001 |
| Cocaine use |  |  | 3.19 (1.17-8.74) | 0.024 |
| Alcohol use |  |  | 0.29 (0.065-1.31) | 0.11 |
| Opioid use | 3.22 (1.89-5.65) | <0.001 | 2.48(1.40-4.39) | 0.002 |
| Non-adherence | 1.81 (1.02-3.19) | 0.042 | 1.06 (1.04-1.08) | <0.001 |
| Total SLEDAI-2K |  |  | 1.60 (0.90, 2.85) | 0.11 |

First row is a model with opioid use and SLE disease activity (SLEDAI-2K)
Second row is a model with opioid use and alcohol use,
Third row is a model with opioid use, cocaine use and alcohol use
Fourth row is opioid use, non-adherence, and SLEDAI-2K.
The models demonstrate opioid use and non-adherence with medical therapy are the only consistent predictors, and that non-adherence dominates over SLE disease activity (SLEDAI-2K) in the last model.

The Multivariate Cox Model was used to estimate for probability of death in SLE patients. Using covariates opioid use and total SLEDAI-2K, they were both statistically significant, with hazard ratios of 2.59 and 1.06 respectively (Table 3).

Using covariates opioid use and alcohol use, only opioid use was statistically significant with a hazard ratio of 3.28 (Table 3).

Using covariates opioid use, cocaine use, and alcohol use, only opioid use and cocaine use were statistically significant, with hazard ratios of 2.99 and 3.19 respectively (Table 3).

Lastly, using covariates opioid use, non-adherence, and total SLEDAI-2K, only opioid use and non-adherence were statistically significant, with hazard ratios 2.48 and 1.06 respectively (Table 3).

The marginal survival for lupus patients not taking opioids was 87.6% (12.4% dead) versus 65.2% (34.8% dead) patients taking opoiods (p<0.001). The Kaplan-Meir survival curve revealed higher probability of survival for lupus patients that did not use opioids (p<0.001) (Figure 1). The Kaplan-Meir survival curve is telling in that most of the deaths occur in the first 5 years of disease in both groups and then the survival stabilized. Thus, this structure to the curve suggests that opioid use is most risky to those patients with recently diagnosed disease, rather than those with chronic disease who seem to have accomodated to chronic opioid use.

**Figure 1.**
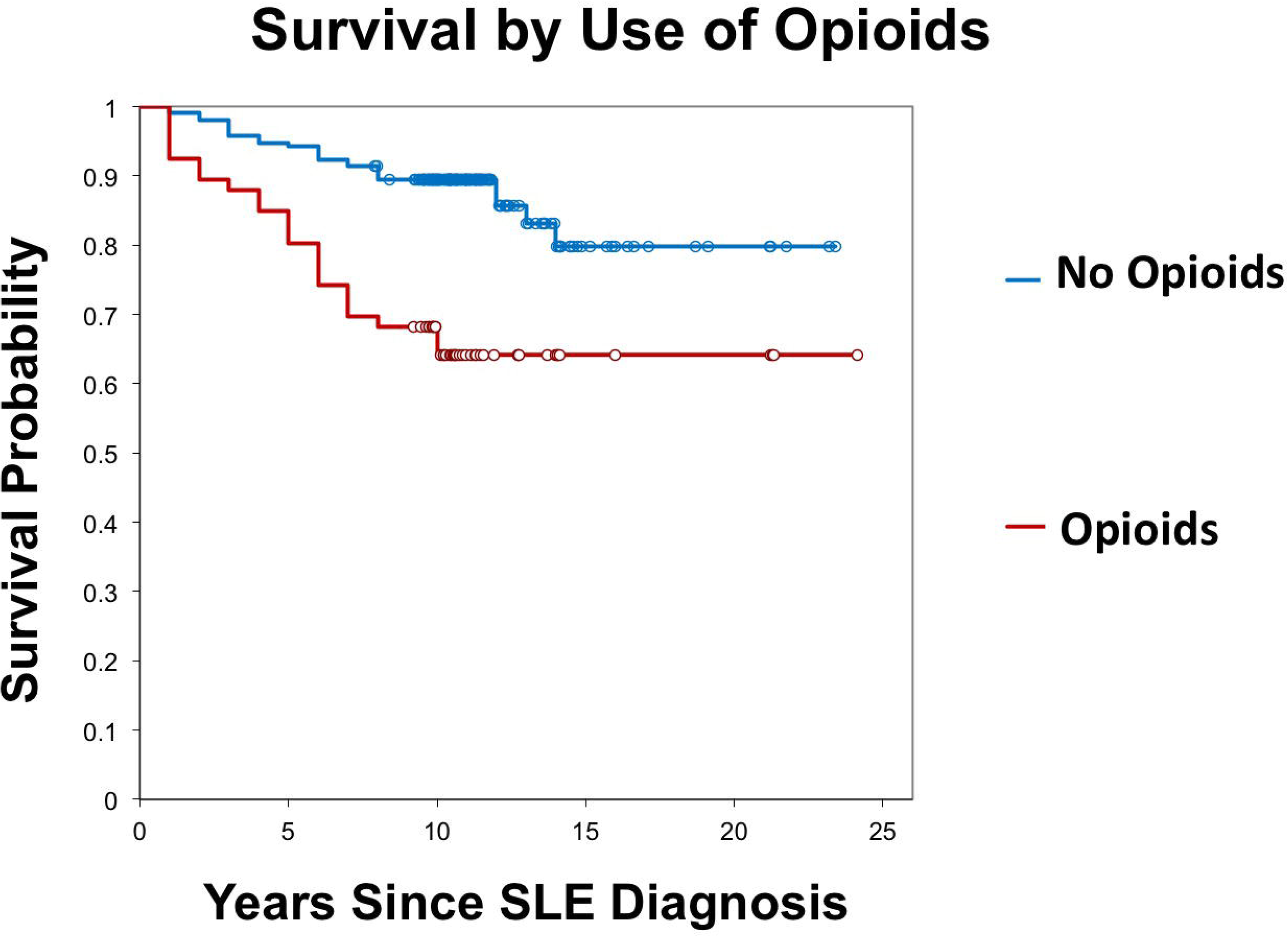
The Kaplan-Meir survival curve showing a higher probability of survival for SLE patients who did not use opioids (p<0.001).

## Discussion

In the last few decades, a general consensus arose that cancer pain and acute post-surgical pain were being undertreated because of physician fear of the toxicity and addictive qualities of opioid analgesics, and subsequently the Agency for Healthcare Research and Quality created a number of guidelines for the assessment of pain using metrics and management of acute pain in a number of clinical situations, recommendations that led routine pain assessment and to a greater use of opioids in the acute care setting (23,24). Although certain guidelines recommended narcotic analgesics to manage recalcitrant chronic pain in osteoarthritis and rheumatoid arthritis, other guidelines minimized the use of opioids and recommended other non-opioid methods of pain control (25,26). However, opioid analgesic use has markedly increased among arthritis patients followed by numerous reports of adverse opioid effects on outcome in certain populations (8,9,10,27-32).

SLE is often characterized by substantial chronic pain related to arthritis, neuropathy, Raynaud’s phenomenon, low back pain, vertebral compression fractures, headache, fibromyalgia and aseptic necrosis of the hips (1-5). A substantial proportion of SLE patients (up to 42%) show signs of pain-related dysfunction, including distress, activity interference, catastrophising, and interpersonal difficulties (2,3,33-35). Treating chronic pain in these individuals is rather difficult due to concomitant coagulation and renal disorders that limit nonsteroidal anti-inflammatory drug use, thus, there has been impetus to treat certain types of chronic pain in SLE with opioids in a parallel fashion to the opioid analgesic approach to rheumatoid arthritis in order to avoid the potential toxic effects of non-steroidal anti-inflammatory drugs and excessive corticosteroids (6-8).

The use of opioid analgesics in rheumatoid arthritis for chronic pain has been associated with increased serious adverse events including safety events, falls, fractures and increased all-cause mortality (9,10). The increased mortality associated with opioid analgesics appears to be in part due to increased accidents and unobserved cardiopulmonary arrest (31,32,36). Recently Krebs and colleagues have demonstrated in a randomized trial that treatment with opioids is not superior to treatment with non-opioid medications for improving pain-related function of chronic low back pain and hip or knee osteoarthritis, supporting the use of non-opioid approaches for managing chronic musculoskeletal pain (37).

The present study demonstrates that chronic opioid analgesic use in SLE is common (24% prevalence) and is associated with an independent mortality risk that in the multivariate model cannot be statistically accounted for by disease activity, disease injury, or other SLE-related factors (Tables 2 and 3). The causes of opioid-related deaths are multifactorial, including prescriber behaviors, patient contributory factors, disease-related factors, nonmedical use patterns, and systemic failures, and all have to be addressed in a root-analysis approach (36).

Many of the deaths that occurred in the present study were unobserved deaths outside of the hospital similar to the recognized pattern of non-adherence to medical therapy, drug-induced accidents, unintentional death, and cardiopulmonary arrest (36). Autopsy series of SLE deaths have typically demonstrated a number of characteristic SLE-related pathologic findings but the actual terminal event in many patients could not be identified and thus, non-disease related factors are often implicated in the terminal event (38-41). There is a similar increase in mortality in renal transplant patients who utilize chronic opioid analgesics, and this might be due to associated disease severity rather than to direct opioid effects (42). This present study indicates that opioid use in SLE is indeed associated with higher total SLICC/ACRDI and total SLEDAI-2K scores indicating more severe and active disease in opioid users, thus, opioid use being an marker of disease severity or non-adherence to therapy may also be correct in SLE populations (42).

Substance abuse in SLE patient is not common, but alcohol abuse, cocaine use, nicotine addiction, and opioid addiction have been reported (43-47). SLE patients may also have a number of psychological complaints including depression, anxiety, amnestic complaints, disordered thinking, concentration deficits, as well as frank neurologic disorders including seizures, epilepsy, encephalopathy, stroke, and cord syndromes amongst others (18,35,48). A certain number of these conditions co-exist with superimposed pain-related dysfunction, including distress, activity interference, catastrophising, and interpersonal difficulties, and these can be complicated by effects of opioid analgesics (2-3,33-35). Suicide risk is also increased in SLE, and suicide can be associated with excessive opioid use (49-50). However, it is most likely that opioid-related deaths in SLE are not suicide, but rather unintentional, related to accidents, falls, cardiopulmonary arrest, concomitant use of tobacco or cocaine, non-adherence to medical therapy, or associated disease severity (31,32,36,42).

There are a number of limitations to this study. The period of observation was 5 years, longer or shorter periods of observation may have different results. The present study was a cross-sectional, prospective cohort study that accurately reflected our SLE population over 5 years and provided opioid use prevalence figures, but was not a randomized controlled trial of opioid verses non-opioid analgesics, thus, the reported associations may not be causative (37). The present study was not an inception cohort, thus, there may be bias related to over-inclusion of survivors with long-term disease and exclusion of individuals with severe disease who had already died before inclusion. However, this type of bias would tend to lessen the association of mortality with opioid use and was controlled for to some extent by excluding severely ill hospitalized SLE patients, and in the statistical analysis cofactoring for disease activity, disease injury, concomitant substance abuse, and non-adherence to medical therapy. “Missingness” is a problem in many mortality studies; however, a strength of this study was the minimization of “missingness” by using a combination of local residence requirement for enrollment, extended family and friends contacts, the exclusion of individuals without governmental identification, the exclusion of acutely ill hospitalized patients, the use of autopsy files from the medical examiner, and the search for missing subjects by the examination of medical records, newspaper obituary databases, and confirmatory governmental death and benefit indexes.

## Conclusion

This study indicates that chronic opioid analgesic use is common (24%) in SLE patients. Opioid analgesic use in SLE is associated with more severe and active disease, increased prevalence of tobacco and cocaine use, non-adherence to medical therapy, and increased mortality. In each statistical model, opioid use was a strong and independent risk factor for mortality in SLE. The increased mortality in the opioid analgesic cohort might not be a direct opioid effect, but rather related to associated increased SLE disease injury, increased SLE disease activity, non-adherence to therapy, concomitant tobacco and cocaine use, or a combination of these or other factors. Nevertheless, these findings suggest that opioid analgesics should be used cautiously in SLE patients, and emphasize the need for appropriate patient education, safe opioid-prescribing practices, and alternative non-opioid strategies for managing chronic pain.

Disclosure Statement: None of the authors declare a conflict of interest.

## REFERENCES

1. Mahmoud K, Zayat A, Vital EM. Musculoskeletal manifestations of systemic lupus erythmatosus. Curr Opin Rheumatol 2017;29:486–492.

2. Greco, C. M., Rudy, T. E., & Manzi, S. Adaptation to Chronic Pain in Systemic Lupus Erythematosus: Applicability of the Multidimensional Pain Inventory. Pain Medicine 2003;4:39– 50.

3. Özel F, Argon G. The effects of fatigue and pain on daily life activities in systemic lupus erythematosus. Agri. 2015;27:181–9.

4. Di Franco M, Guzzo MP, Spinelli FR, Atzeni F, Sarzi-Puttini P, Conti F, Iannuccelli C. Pain and systemic lupus erythematosus. Reumatismo 2014;66:33–8.

5. Iannuccelli C, Spinelli FR, Guzzo MP, Priori R, Conti F, Ceccarelli F, Pietropaolo M, Olivieri M, Minniti A, Alessandri C, Gattamelata A, Valesini G, Di Franco M. Fatigue and widespread pain in systemic lupus erythematosus and Sjögren’s syndrome: symptoms of the inflammatory disease or associated fibromyalgia? Clin Exp Rheumatol. 2012;30(6 Suppl 74):117–21.

6. Borenstein DG, Hassett AL, Pisetsky D. Pain management in rheumatology research, training, and practice. Clin Exp Rheumatol. 2017 Sep-Oct;35 Suppl 107(5):2–7. Epub 2017 Sep 28. Review. PMID:28967362

7. Curtis JR, Xie F, Smith C, Saag KG, Chen L, Beukelman T, Mannion M, Yun H, Kertesz S. Changing trends in opioid use among patients with rheumatoid arthritis in the United States. Arthritis Rheumatol 2017;69:1733–1740.

8. Zamora-Legoff JA, Achenbach SJ, Crowson CS, Krause ML, Davis JM 3rd, Matteson EL. Opioid use in patients with rheumatoid arthritis 2005-2014: a population-based comparative study. Clin Rheumatol 2016;35:1137–44.

9. Shipton EA, Shipton EE, Shipton AJ. A Review of the Opioid Epidemic: What Do We Do About It? Pain Ther. 2018 Apr 6. doi: 10.1007/s40122-018-0096-7. [Epub ahead of print] Review. PMID: 29623667

10. Lo-Ciganic WH, Floden L, Lee JK, Ashbeck EL, Zhou L, Chinthammit C, Purdy AW, Kwoh CK. Analgesic use and risk of recurrent falls in participants with or at risk of knee osteoarthritis: data from the Osteoarthritis Initiative. Osteoarthritis Cartilage. 2017 Sep;25(9):1390–1398. doi: 10.1016/j.joca.2017.03.017. Epub 2017 Apr 4. PMID: 28385483

11. Guy GP Jr, Zhang K, Bohm MK, Losby J, Lewis B, Young R, Murphy LB, Dowell D. Vital Signs: Changes in opioid prescribing in the United States, 2006-2015. MMWR Morb Mortal Wkly Rep. 2017; 66:697–704.

12. Tan EM, Cohen AS, Fries JF, Masi AT, McShane DJ, Rothfield NF, et al. 1982 Revised criteria for the classification of systemic lupus erythematosus. Arth Rheum 1982;25:1271–1277.

13. Hochberg MC. Updating the American College of Rheumatology revised criteria for the classification of systemic lupus erythematosus. Arthritis Rheum 1997;40:1725–26.

14. Gladman DD, Ibañez D, Urowitz MB. Systemic lupus erythematosus disease activity index 2000. J Rheumatol. 2002 Feb;29(2):288–91.

15. Gladman D, Ginzler E, Goldsmith C, Fortin P, Liang M, Urowitz M, et al. The development and initial validation of the Systemic Lupus International Collaborating Clinics/American College of Rheumatology damage index for systemic lupus erythematosus. Arthritis Rheum 1996;39:363–369.

16. Sibbitt WL Jr, Brandt JR, Johnson CR, Maldonado ME, Patel SR, Ford CC, Bankhurst, AD, Brooks WM: The incidence and prevalence of neuropsychiatric syndromes in pediatric-onset systemic lupus erythematosus. J Rheum 2002;29:1536–42.

17. Ghaussy NO, Sibbitt W Jr, Bankhurst AD, Qualls CR: The effect of race on disease activity in systemic lupus erythematosus. J Rheumatol 2004;31:915–9.

18. American College of Rheumatology. The American College of Rheumatology nomenclature and case definitions for neuropsychiatric lupus syndromes. Arthritis Rheum 1999;42:599–608.

19. Centers for Disease Control and Prevention. CDC Guideline for Prescribing Opioids for Chronic Pain. https://www.cdc.gov/drugoverdose/prescribing/guideline.html accessed 09/29/2017.

20. Katz J, Melzack R. Measurement of pain. Surg Clin North Am. 1999;79:231–252.

21. Myles PS, Troedel S, Boquest M, Reeves M. The pain visual analogue scale. Is it linear or nonlinear? Anesth Analg 1999;89:1517–20.

22. U.S., Social Security Death Index, 1935-. Ancestry.com. Original data: Social Security Administration. Social Security Death Index, Master File. Social Security Administration, USA. http://search.ancestry.com/search/db.aspx?dbid=3693, Accessed Sept.25, 2017.

23. Steele JA, Taylor E. Guidelines released on acute pain management. J Natl Cancer Inst 1992;84:481–2.

24. McCann J. New guidelines may unify cancer pain management. J Natl Cancer Inst 1998;90:733–4

25. Vasudevan SV, Potts EE, Mehrotra C. Pain management in arthritis: evidence-based guidelines. WMJ 2003;102:14–8.

26. American College of Rheumatology. Recommendations for the medical management of osteoarthritis of the hip and knee: 2000 update. American College of Rheumatology Subcommittee on Osteoarthritis Guidelines. Arthritis Rheum. 2000;43:1905–15.

27. Clausen, T., Waal, H., Thoresen, M., & Gossop, M. Mortality among opiate users: opioid maintenance therapy, age and causes of death. Addiction 2009;104:1356–1362.

28. Gomes, T, Mamdani, MM, Dhalla, IA, Paterson, JM. Juurlink, DN. Opioid dose and drug-related mortality in patients with nonmalignant pain. Arch of Intern Med 2011;171:686–91

29. Manchikanti, L, Singh, A. Therapeutic opioids: a ten-year perspective on the complexities and complications of the escalating use, abuse, and nonmedical use of opioids. Pain Physician 2008;11(Suppl 2),63–88.

30. Miller, NS: Failure of enforcement of controlled substance laws in health policy for prescribing opiate medications: a painful assessment of morbidity and mortality. Am J Ther 2006;13:527–33.

31. Paulozzi LJ, Ryan GW. Opioid analgesics and rates of fatal drug poisoning in the United States. Am J Prev Med. 2006;31:506–11.

32. Paulozzi LJ, Budnitz DS, Xi Y. Increasing deaths from opioid analgesics in the United States. Pharmacoepidemiol Drug Saf. 2006 Sep;15(9):618–27.

33. Fischin J, Chehab G, Richter JG, Fischer-Betz R, Winkler-Rohlfing B, Willers R, Schneider M. Factors associated with pain coping and catastrophising in patients with systemic lupus erythematosus: a cross-sectional study of the LuLa-cohort. Lupus Sci Med 2015;2:e000113. doi: 10.1136/lupus-2015-000113. eCollection 2015

34. Somers TJ, Kurakula PC, Criscione-Schreiber L, Keefe FJ, Clowse ME. Self-efficacy and pain catastrophizing in systemic lupus erythematosus: relationship to pain, stiffness, fatigue, and psychological distress. Arthritis Care Res (Hoboken). 2012;64:1334–40.

35. Kozora E, Ellison MC, West S. Depression, fatigue, and pain in systemic lupus erythematosus (SLE): relationship to the American College of Rheumatology SLE neuropsychological battery. Arthritis Rheum. 2006;55:628–35.

36. Webster LR, Cochella S, Dasgupta N, Fakata KL, Fine PG, Fishman SM, Grey T, Johnson EM, Lee LK, Passik SD, Peppin J, Porucznik CA, Ray A, Schnoll SH, Stieg RL, Wakeland W. An analysis of the root causes for opioid-related overdose deaths in the United States. Pain Med. 2011;12 Suppl 2:S26–35.

37. Krebs EE, Gravely A, Nugent S, Jensen AC, DeRonne B, Goldsmith ES, Kroenke K, Bair MJ, Noorbaloochi S. Effect of Opioid vs Nonopioid Medications on Pain-Related Function in Patients With Chronic Back Pain or Hip or Knee Osteoarthritis Pain: The SPACE Randomized Clinical Trial. JAMA. 2018 Mar 6;319(9):872–882.

38. Sibbitt WL Jr., Brooks WM, Kornfield M, Hart BL, Bankhurst AD, Roldan CA: Magnetic resonance imaging and brain histopathology in neurospyschiatric systemic lupus erythematosus. Semin Arthritis Rheum. 2010;40:32–52.

39. Ellis SG,Verity MA. Central nervous systemic involvement in systemic lupus erythematosus: a review of neuropathological findings in 57 cases, 1955-1977. Semin Arthritis Rheum 1979;8:212–221.

40. Hanly JG, Walsh N, Sangalang V. Brain pathology in systemic lupus erythematosus J Rheumatol 1992;19:732–741.

41. Devinsky O, Petito CK, Alonso DR. Clinical and neuropathological findings in systemic lupus erythematosus: The role of vasculitis, heart emboli, and thrombotic thrombocytopenic purpura. Ann Neurol 1988;23:380–384.

42. Abbott KC, Fwu CW, Eggers PW, Eggers AW, Kline PP, Kimmel PL. Opioid prescription, morbidity, and mortality in US transplant recipients. Transplantation. 2018 Jan 10. doi: 10.1097/TP.0000000000002057. [Epub ahead of print] PMID: 29319627

43. Gerevich J. Compulsive heroin use: comorbidity, syndrome or self-medication of lupus erythematosus? Acta Psychiatr Scand. 2001;103:78–9.

44. Gerevich J, Bácskai E, Farkas L. Use of heroin to cope with stress caused by a negative life event in a patient with lupus erythematosus. Psychosom Med. 2005;67:341

45. van Weelden M, Queiroz LB, Lourenço DM, Kozu K, Lourenço B, Silva CA. Alcohol, smoking and illicit drug use in pediatric systemic lupus erythematosus patients. Rev Bras Reumatol Engl Ed 2016;56:228–34.

46. Takvorian SU, Merola JF, Costenbader KH. Cigarette smoking, alcohol consumption and risk of systemic lupus erythematosus. Lupus. 2014;23:537–44.

47. Ghaussy NO, Sibbitt WL Jr, Qualls CR. Cigarette smoking, alcohol consumption, and the risk of systemic lupus erythematosus: a case-control study. J Rheumatol. 2001;28:2449–53.

48. Peralta-Ramírez MI, Jiménez-Alonso J, Godoy-García JF, Pérez-García M; Group Lupus Virgen de las Nieves. The effects of daily stress and stressful life events on the clinical symptomatology of patients with lupus erythematosus. Psychosom Med. 2004;66:788– 94.

49. Matsukawa Y. Suicide in systemic lupus erythematosus. Psychosomatics. 2015;56:317–8.

50. Roy A. Characteristics of opiate dependent patients who attempt suicide. J Clin Psychiatry. 2002;63:403–7.

